# Wireless networks of injectable microelectronic stimulators based on rectification of volume conducted high frequency currents

**DOI:** 10.1101/2022.03.11.483920

**Authors:** Aracelys García-Moreno, Albert Comerma-Montells, Marc Tudela-Pi, Jesus Minguillon, Laura Becerra-Fajardo, Antoni Ivorra

## Abstract

**Objective:** To develop and *in vivo* demonstrate threadlike wireless implantable neuromuscular microstimulators that are digitally addressable.

**Approach:** These devices perform, through its two electrodes, electronic rectification of innocuous high frequency current bursts delivered by volume conduction via epidermal textile electrodes. By avoiding the need of large components to obtain electrical energy, this approach allows the development of thin devices that can be intramuscularly implanted by minimally invasive procedures such as injection. For compliance with electrical safety standards, this approach requires a minimum distance, in the order of millimeters or a very few centimeters, between the implant electrodes. Additionally, the devices must cause minimal mechanical damage to tissues, avoid dislocation and be adequate for long-term implantation. Considering these requirements, the implants were conceived as tubular and flexible devices with two electrodes at opposite ends and, at the middle section, a hermetic metallic capsule housing the electronics.

**Main results:** The developed implants have a submillimetric diameter (0.97 mm diameter, 35 mm length) and consist of a microcircuit, which contains a single custom-developed integrated circuit, housed within a titanium capsule (0.7 mm diameter, 6.5 mm length), and two platinum-iridium coils that form two electrodes (3 mm length) located at opposite ends of a silicone body. These neuromuscular stimulators are addressable, allowing to establish a network of microstimulators that can be controlled independently. Their operation was demonstrated by injecting a few of them in the hind limb of anesthetized rabbits and inducing controlled and independent contractions.

**Significance:** These results show the feasibility of manufacturing threadlike wireless addressable neuromuscular stimulators by using fabrication techniques and materials well established for chronic electronic implants. This paves the way to the clinical development of advanced motor neuroprostheses formed by dense networks of such wireless devices.

## 1. Introduction

Most implantable neuroprosthetic systems based on electrical stimulation consist of an electronic unit, typically referred to as implantable pulse generator (IPG), wired to electrodes at the target sites (e.g., nerves and muscles). This approach has been successfully adopted for developing sensory neuroprostheses, such as cochlear implants for the deaf [1], for developing motor neuroprostheses, such as drop foot stimulators for improving gait in patients with hemiplegia [2], and for epidural electrical stimulators for restoring walking after paralysis [3]. These successful instances correspond either to situations in which a single target must be electrically stimulated, or to situations in which multiple targets must be stimulated but these are clustered in a small and relatively static region. Such situations allow establishing the electrical connections between the IPG and the electrodes with a single or a few multiwire cables. There are also instances of clinically demonstrated motor neuroprostheses with a few targets distributed over relatively large and mobile regions [4,5]. However, this wired and centralized topology is generally deemed as problematic, particularly for developing motor neuroprostheses, because it involves highly invasive surgeries and frequently results in mechanical failure of electrical connections [6]. As a consequence, few targets are independently targeted in wired motor neuroprostheses, thus limiting the potential of multi-channel control.

In 1991, G.E. Loeb and colleagues introduced a novel paradigm: they conceived and developed capsules that integrated both the IPG and the electrodes and that were small enough for being intramuscularly delivered by percutaneous injection at multiple target sites [7]. These devices were later referred to as BIONs and, having 256 independent addresses, were in principle capable of forming a dense network of single-channel wireless microstimulators for finely coordinated neuromuscular stimulation. The original, and smallest, glass BION devices had a length of about 16 mm and a diameter of 2 mm and were energized solely by inductive coupling with an external coil to be worn by the patient. Later ceramic devices overcame fragility issues in the original glass capsules and contained a rechargeable battery for discontinuous use of the external coil. These enhancements came at the cost of a larger size: newer BIONs had a length of 27 mm and a diameter of 3.3 mm [6]. BIONs were successfully employed for treatment of some ailments such as poststroke shoulder subluxation in hemiplegic subjects [8]. More recently, intramuscularly implantable myographic sensors based on the BION technology have been demonstrated for controlling robotic arms in amputees [9]. However, despite their successes, BIONs are too bulky for massive implantation and the dense network of microstimulators concept was never implemented.

Further miniaturization is hindered in BION-like devices – and in electronic medical implants in general – due to the need of bulky components to provide electrical energy.

Wireless power transfer (WPT) is frequently used in electronic medical implants as an alternative to electrochemical batteries. WPT offers two major benefits: implant longevity and miniaturization.

To the best of our knowledge, the only WPT methods in clinical use are near-field inductive coupling and ultrasonic acoustic coupling [10], the former being much more prevalent than the latter. Other WPT methods under exploration are: optic WPT [11–15], mid-field inductive coupling [16,17], far-field coupling [18], capacitive coupling [19–21] and WPT based on volume conduction, which is also, less accurately, referred to as galvanic coupling [22–25]. Comprehensive and partial recent reviews on these methods can be found in [26–29].

Compared to other WPT methods, galvanic coupling and capacitive coupling offer the advantage of not integrating bulky parts, such as piezoelectric crystals or coils, within the implant for receiving the energy transferred by the remote transmitter. As demonstrated in [30], the energy can be readily picked up with a pair of thin electrodes separated by a few millimeters or centimeters.

Recently, it has been proposed the development of microstimulators whose operation is based on rectification of volume conducted innocuous high frequency (HF) current bursts [31]. By passively rectifying HF currents across two electrodes, the devices cause local low frequency (LF) currents capable of stimulation of excitable tissues without requiring large components such as coils or batteries. A portion of the HF current picked up by the devices, instead of being rectified for stimulation, can be used to power digital electronics (e.g., for controlling the rectification process and for communications) and, in this sense, it can be understood that the proposed devices employ WPT based on volume conduction. The current bursts, at frequencies between 1 and 10 MHz, are delivered by an external generator and are supplied to the tissues through epidermal electrodes.

The proposed method offers the possibility to build small devices that can be implanted intramuscularly by minimally invasive procedures, such as percutaneous injection, thus reducing damage to tissues.

As analyzed in [32], for ensuring compliance with electrical safety standards, WPT based on HF volume conduction requires a minimum distance, in the order of millimeters or a very few centimeters, between the implant electrodes to obtain enough power for electronic operation and electrical stimulation of tissues. Additionally, the device itself must cause minimal damage to tissues, avoid implant dislocation and be adequate for long-term implantation. For these reasons, the implants, according to the proposed method, were conceived as tubular, elongated, flexible and thin devices with two electrodes at opposite ends.

Proof-of-concept prototypes fulfilling these characteristics have been previously developed and *in vivo* demonstrated. In [33], flexible threadlike injectable microstimulators – with a diameter of 1 mm and a length of 3 cm, and composed only of a diode, two capacitors, and a resistor – were capable of performing non-addressable charge-balanced intramuscular stimulation. Addressability was later demonstrated in [22], where 2 mm thick semi-rigid microcontroller-based injectable stimulators were able to perform controlled intramuscular stimulation of antagonist muscles in an independent manner by digitally addressing them.

Nevertheless, aiming at clinical use, more advanced prototypes need to be demonstrated. In particular, it must be demonstrated that the proposed method is compatible with materials and construction techniques that are deemed as adequate by regulatory bodies to ensure biocompatibility and long-term operation. In this regard, one aspect is particularly challenging: the active electronics of the implant must be hermetically housed to ensure long-term operation and, concurrently, the hermetic housing must not hamper minimal invasiveness.

Here we present the development and the *in vivo* demonstration of very thin flexible injectable microstimulators that contain a metallic capsule housing a hybrid microcircuit mainly formed by a single Application Specific Integrated Circuit (ASIC). These threadlike implants, with a submillimetric diameter, are addressable, allowing to establish a dense network of microstimulators that can be controlled independently (**Figure 1**).

**Figure 1.**
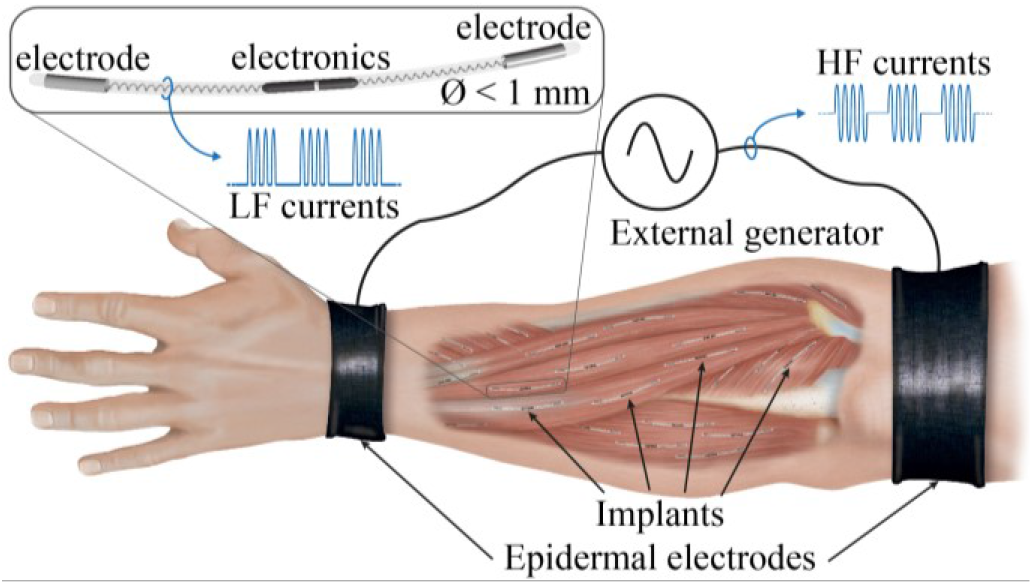
The addressable neuromuscular microstimulators demonstrated here are based on inducing low frequency (LF) currents by electronic rectification of high frequency (HF) bursts delivered via volume conduction through epidermal electrodes. We envision the development of advanced motor neuroprostheses formed by dense networks of such wireless devices which will be implanted by injection.

## 2. Methods

### 2.1 Implant design and development

The injectable microstimulators consist of a very thin, flexible, and elongated tubular body (0.97 mm diameter, 35 mm length). The body, mostly made of silicone, contains in its middle a titanium capsule (0.7 mm diameter, 6.5 mm length) housing the electronic circuit capable of charge-balanced rectification of HF currents delivered by volume conduction. Two platinum-iridium (Pt-Ir) helicoidal coils with variable pitch form the two electrodes (3 mm length), located at opposite ends of the device, and the interconnections from these two electrodes to the two feedthroughs at opposite ends of the capsule (**Figure 2**).

**Figure 2.**
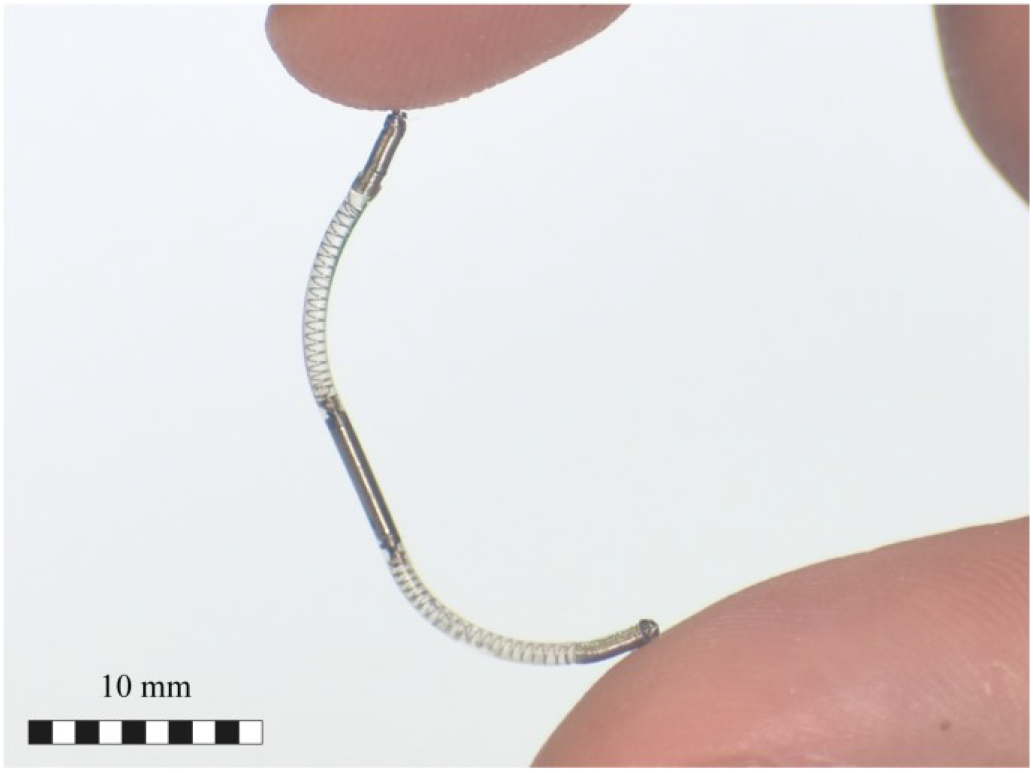
Picture of the implemented device with a diameter of 0.97 mm and a length of 35 mm.

#### 2.1.1 Implant circuit

The circuit of the implants consists of an ASIC and three capacitors (**Figure 3**). In essence, the integrated circuit acts as a controllable rectifier. Through the implant electrodes, the circuit collects HF current during the bursts. A portion of such HF current is used to power a digital circuit. The digital circuit, in response to commands from the external system modulated within the bursts, dictates the rectification of HF current through the electrodes thereby generating LF currents capable of stimulation.

**Figure 3.**
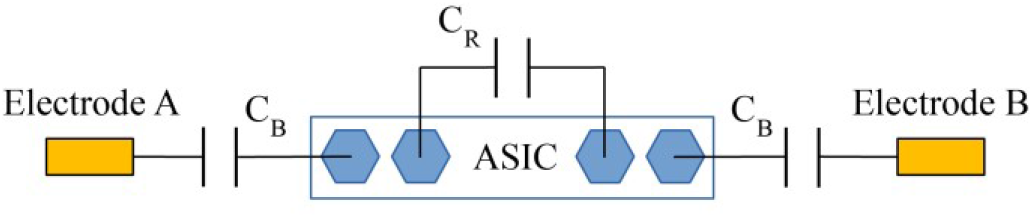
General circuit architecture of the addressable microstimulator capable of performing charge-balanced intramuscular stimulation. C_B_ are the dc-blocking capacitors and C_R_ is the power supply reservoir capacitor.

Two dc-blocking capacitors (C_B_) are connected in series at each input of the ASIC to prevent any flow of dc current through the implant electrodes, and a power supply reservoir capacitor (C_R_) is connected to store the necessary charge to provide continued power to the device in between bursts. Due to their relatively large capacitance values (1 μF and 0.22 μF), these capacitors cannot be integrated in the ASIC without greatly increasing its size and hence were mounted externally.

It is worth noting that two dc-blocking capacitors were mounted, instead of just a single capacitor, for maximizing the combined voltage rating (6.3 V + 6.3V) and for achieving single fault safety.

#### 2.1.2 ASIC circuit architecture

The HF bursts are amplitude modulated to enable selecting which device will produce stimulation, using an internally defined address, and to enable selecting the amplitude of the stimulation current.

The simulation of the stimulation sequence depicted in **Figure 4** facilitates understanding of the operation of the ASIC. A Thévenin equivalent, consisting of a voltage source (V_Th_) and a series resistance (R_Th_), models the presence of the HF electric field at the location of the implant. The resistance (R_Th_) approximates the tissue impedance magnitude across the implant electrodes at the desired operating frequency (*f*) as in [24].

**Figure 4.**
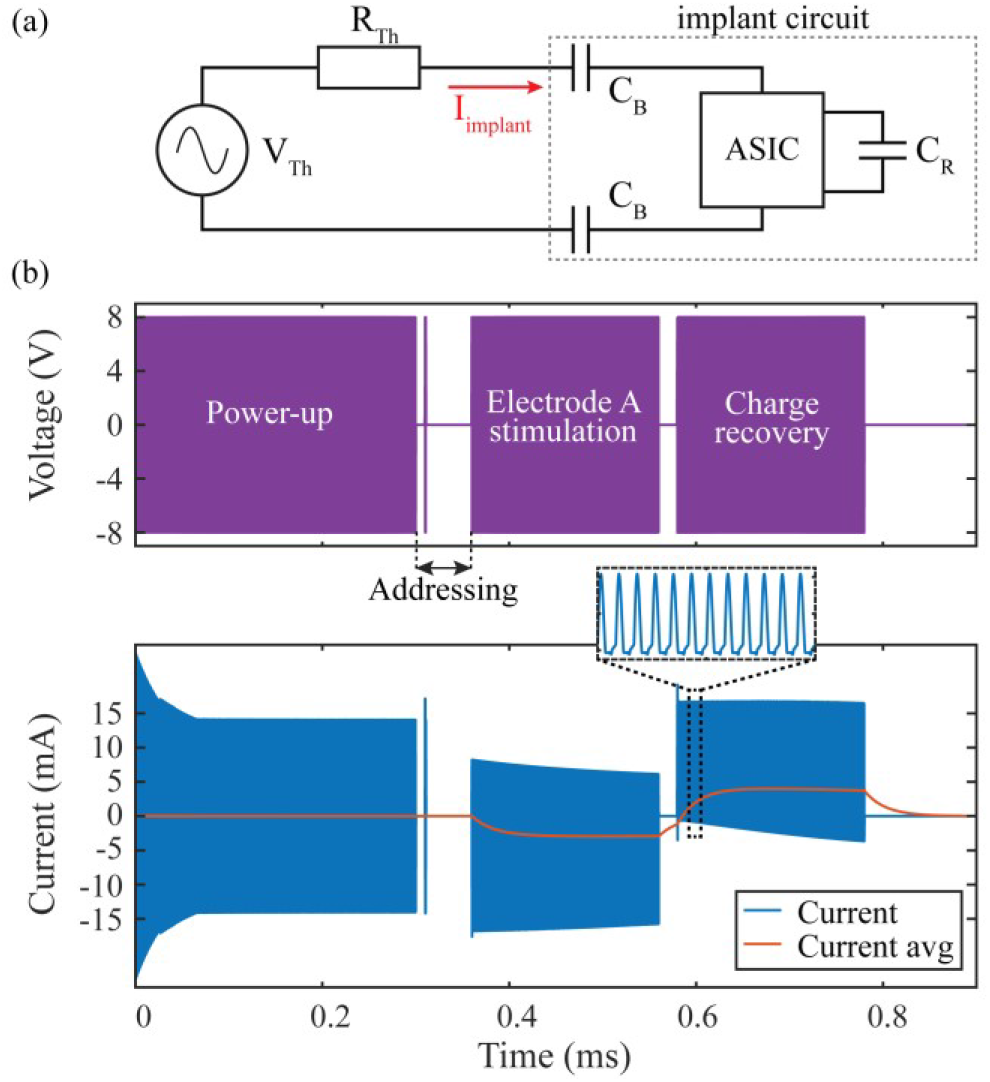
Simulation of the ASIC addressing and stimulation sequence at an operating frequency of 3 MHz: (a) Thévenin equivalent model of the HF electric field at implant location. (b) Voltage from external generator (purple), implant current (blue) and low frequency current (red) as obtained by filtering the implant current with a first-order low-pass filter with a cut-off frequency of 7.2 kHz.

Each modulated HF burst consists of a sequence of sub-bursts. The stages of a stimulation sequence are the following:

- Power-up: A long sub-burst (300 μs) enables charging the reservoir capacitor (C_R_) and initiates several internal subcircuits of the ASIC.
- Addressing: during 64 μs the ASIC monitors the voltage to detect the presence of a very-short sub-burst that determines whether the device must be enabled (the time position of the sub-burst indicates the address).
- Current setting: the presence of a second very-short sub-burst at the end of the 64 μs interlude determines whether the magnitude of the stimulation current is 4 mA (by default 2 mA). (An example of magnitude current setting at 4 mA is seen in **Figure 14**).
- Cathodic stimulation: if the device has been enabled in the addressing stage, the device will perform stimulation (i.e., it will allow passage of rectified current) during the next sub-burst until the external systems stops the sub-burst (the typical duration is 200 μs). Note that in **Figure 4**(b) the LF current through the implant ceases to be zero. As explained in [31], this LF current generated by rectification can cause stimulation of excitatory tissues.
- Gap: between cathodic stimulation and charge recovery, a short gap or delay, also referred to as interphase dwell [34], is introduced for higher stimulation efficacy [35] (the typical duration is 20 μs).
- Charge recovery: in the last stage, if the device has been enabled, passage of rectified current is allowed in the opposite direction to perform active charge recovery (the typical duration is 200 μs).

The ASIC is organized in several subcircuits to provide the intended functionality. **Figure 5** shows the system architecture with the different subcircuits and their main interconnections. In the following paragraphs the function and structure of these subcircuits is overviewed. (A more detailed description will be presented in a subsequent publication).

**Figure 5.**
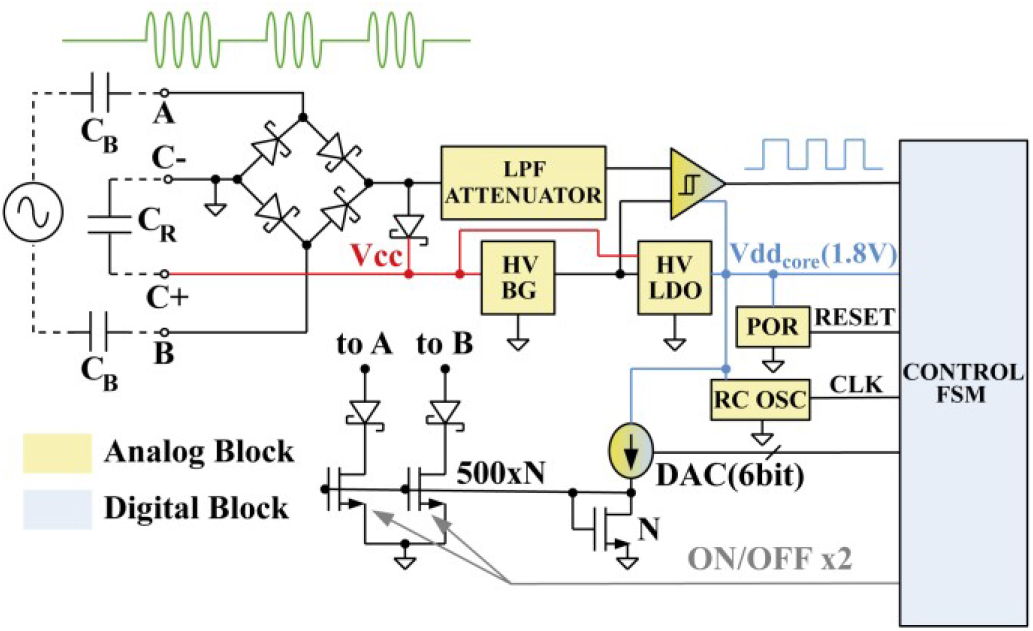
ASIC circuit architecture. The implant electrodes pick up the high frequency current for powering the circuitry through the high voltage (HV) power section consisting of a bridge rectifier, a diode, the external reservoir capacitor (C_R_), a bandgap (BG), a Low Drop Out (LDO) regulator and a comparator. The control finite state machine (FSM) determines whether the device must stimulate and is controlled by a power on reset (POR) and a resistor-capacitance oscillator (RC OSC). The digital to analog converter (DAC) and the current mirror are responsible for controlling the passage of rectified currents for stimulation.

On the HV power side, the fundamental part is the full-wave Schottky rectifier followed by a series diode that feeds the external reservoir capacitor (C_R_). This circuit is the power supply of the system (red node in **Figure 5**). It keeps enough charge stored in the reservoir capacitor to continue powering the device in between the HF sub-bursts. To minimize the capacitance of this capacitor, and hence its geometrical dimensions, it was fundamental to keep the power consumption of the device below 300 μA.

The unregulated voltage generated by the rectifier-capacitor combination is connected to a HV bandgap reference to produce a regulated 1.2 V reference. This voltage together with the unregulated main power is then used by a Low Drop Out (LDO) regulator to power the system with a core voltage of 1.8 V.

A comparator is connected between the rectifier output HV and the unregulated voltage (using resistors to attenuate the voltage) to generate a digital signal for performing communications (i.e., addressing and current control from the external unit). This signal is high when an external sub-burst is present, and it is low when there is no external sub-burst.

It is also important to note the presence of a double HV current mirror connected to the input electrodes through Schottky diodes in series. This subcircuit allows the controlled flow of rectified current and hence it is responsible for performing stimulation. The mirror is very asymmetric to set high currents (in the order of milliamperes) from a relatively small current.

On the core voltage part some subcircuits are included to control the behavior of the digital control finite state machine (FSM). These blocks include a power on reset (POR) to generate a pulse when the power supply rises to the operation voltage level, a relaxation resistor-capacitance (RC) oscillator with a nominal output of 1 MHz, the already mentioned hysteresis comparator to detect the frame signal, and a binary weighted six bit digital to analog converter (DAC) in charge of controlling the output current for stimulation. The main control FSM determines whether it must stimulate or not depending on the incoming frame timing and, if enabled, select the desired output current at the DAC. A voltage to current reference converter is also included to provide a reliable current reference to the DAC.

The design of the ASIC was done in-house using TSMC 0.18 μm Bipolar CMOS DMOS mixed signal, HV process through Europractice. The complete design fits in a very compact form factor of around 300 μm by 2.5 mm. For better integration and reduced volume, the dies provided by the manufacturer with a thickness of approximately 725 μm were thinned down to 8 mils (204 μm).

#### 2.1.3 Electronic assembly

Commercially available ceramic capacitors, with dimensions of 0.6 mm and below, were considered and tested to determine the achievable dimensions of the electronics (**Figure 6**(a)). Two 1 μF capacitors (GRMMDXR60J105ME05 by Murata) were used for dc-blocking (C_B_) and one 0.22 μF capacitor (GRM033R61E224KE01 by Murata) was used as a reservoir capacitor (C_R_) for the internal power supply system.

**Figure 6.**
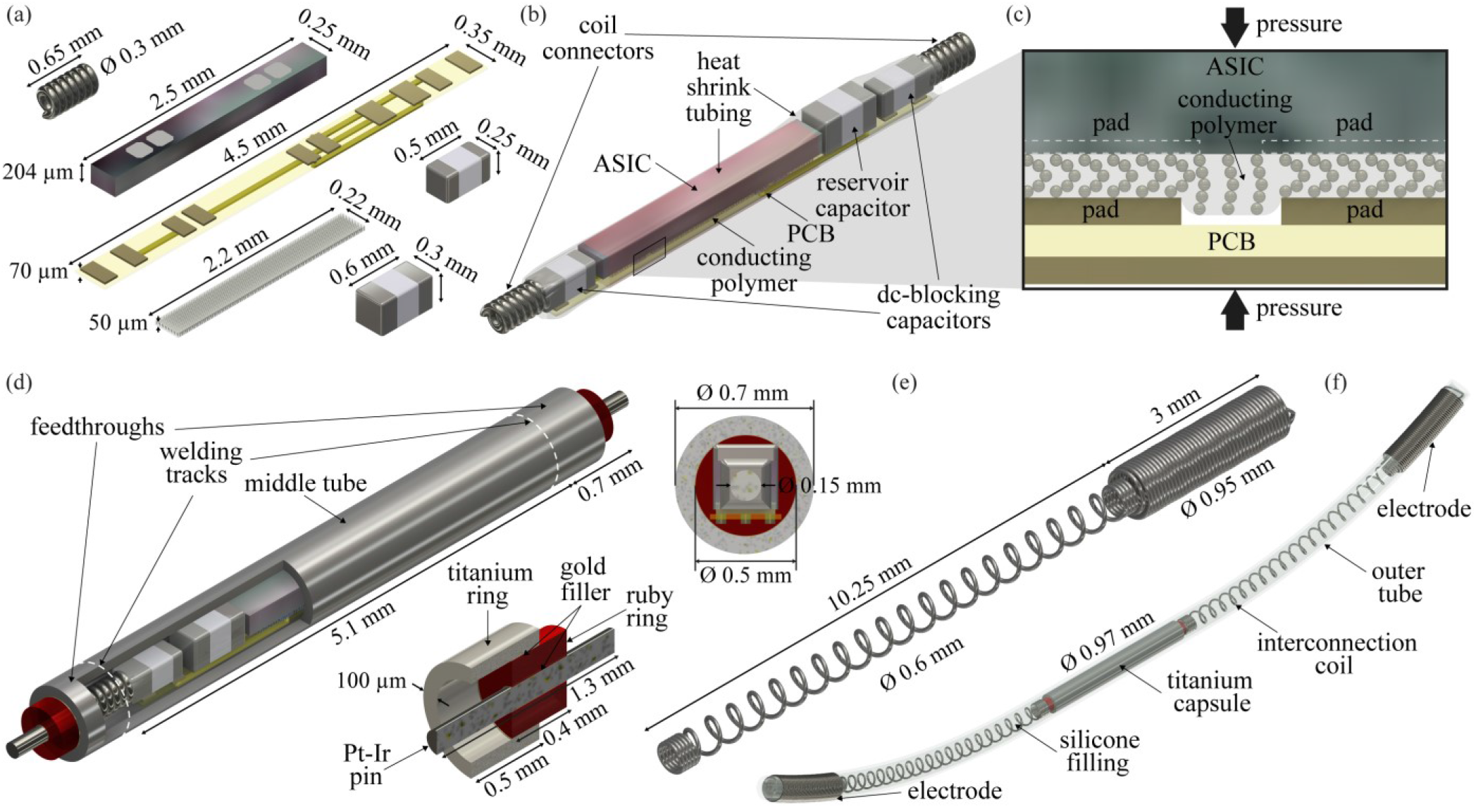
Parts and construction of the injectable microstimulator. (a) Dimensions and (b) assembly of the electronic components. Capacitors are reflow soldered to the printed circuit board (PCB). Two clear ultra-thin medical grade heat shrink tubing are used. The first one (depicted in red for visualization purposes) compresses the conducting polymer and ensures conductivity between ASIC and PCB pads (c). The second one surrounds the assembly for insulation and mechanical robustness. (d) Titanium capsule for the electronic assembly. Two feedthroughs, fabricated using a brazing technique, are laser welded at each end of the middle tube for electrical connections. (e) A Pt-Ir coil with variable pitch and diameter defines the electrode and the interconnection coil at each end of the encapsulated electronics. This coil is laser welded to the feedthrough pin. (f) Final threadlike flexible conformation of the implant. The capsule with the electrodes is inserted in a silicone tube and filled with silicone elastomer, both medical grade, leaving the electrodes exposed and securing the assembly. The implant has a total length of 35 mm, and a maximum diameter of 0.97 mm.

The electronic components were mounted on a 70 μm-thick, 4.5 mm-long double-sided flexible polyimide printed circuit board (PCB) **Figure 6**(a). The capacitors were bonded to the flex PCB using lead-free reflow soldering. The ASIC pads are made of aluminum, which is a non-wettable material, hampering the use of the same method to bond them to the PCB. Traditional methods used for wiring integrated circuits, such as wire bonding, would have substantially increased the volume of the electronic assembly. As an alternative technique to overcome these bonding limitations, it was conceived and tested a novel technique based on the use of conductive polymers and compression force.

The electrical connections between the ASIC and the flex PCB were established placing in between them a 0.05 mm-thick anisotropic conducting polymer with BallWire^®^ technology (PariPoser^®^ by Paricon Technologies Corp.), which provides conductivity in one direction only. These conducting polymers require pressure to ensure minimal resistivity. A clear ultra-thin medical grade thermally shrinkable tubing with a wall thickness of 6 μm and an average diameter of 0.46 mm (103-0335 by Nordson Medical) was placed along the ASIC surrounding the PCB, the conducting polymer and the ASIC, holding these components in place and applying the required pressure (**Figure 6**(b)-(c)).

Pt-Ir wire with a diameter of 0.075 mm was used to manufacture custom-made coiled connectors to facilitate interconnections. Two Pt-Ir microcoils (0.3 mm diameter, 0.65 mm length) soldered to each side of the electronic assembly were used to provide flexibility to the internal connections (**Figure 6**(b)). Overall, the assembly had a maximum diameter of 0.46 mm. A second clear heat shrink tubing with an average diameter of 0.48 mm (103-0782 by Nordson Medical) was applied along the whole assembly surrounding all the electronic components to provide mechanical robustness and electrical insulation without substantially increasing the dimensions of the assembly (**Figure 6**(b)).

#### 2.1.4 Hermetic capsule

Integrated circuitry must be housed hermetically to ensure long-term performance in implants. Silicone rubber – which was used for the encapsulation of the microstimulators developed in [33] and [22] – is a polymer available in medical grade quality and it is frequently used to insulate the electrical cables (i.e., the leads) in long-term active implants in humans due to its high electrical resistivity, its low ion permeability and its versatility and processability. However, silicone rubber, and other polymers, are permeable to water vapor and their use for encapsulating integrated circuitry, in the long-term, leads to corrosion in the delicate microelectronic devices and failure [36,37].

To safeguard the electronics from the harsh environment of the body and simultaneously protect the body from potentially harmful substances, a hermetically sealed encapsulation for the inner electronics was devised. Considering these premises, the dimensions of the electronic assembly, and the mechanical stability required for intramuscular implantation, a titanium capsule and feedthroughs for electrical connections were designed for the encapsulation. The design includes three parts: a middle tube with a wall thickness of 100 μm (inner diameter 0.5 mm, outer diameter 0.7 mm, length 5.1 mm) and two feedthroughs, one at each end. The feedthroughs were fabricated with a ceramic-to-metal sealing technology in which a ruby ring (a stable form of alumina used as the insulating material), Pt-Ir pins (used as electrical conductors) and a titanium ring, were brazed together to achieve high hermeticity (**Figure 6**(d)). As only two electrical connections are required for the assembly, minimum dimensions could be achieved for these parts.

The electronic assembly was gently introduced into the middle tube of the encapsulation, leaving the coiled connectors partially exposed. The feedthrough pins were carefully inserted in the coiled connectors, aligning the three pieces. Holding them together in a custom-made rotation stand, the titanium rings of the feedthroughs were hermetically sealed to the middle tube using a fixed pulsed Nd-YAG laser at a wavelength of 1064 nm in an argon atmosphere (**Figure** 6(d)).

#### 2.1.5 Implant body and electrodes

The conformation and dimensions of the implant were based on those of the implants developed in [33]. The body of the implant was built by filling a silicone tube with uncured silicone. Given that the outer diameter of the hermetic capsule is 0.7 mm, a custom-made silicone tube with a wall thickness of 100 μm (inner diameter 0.72 mm, outer diameter 0.92 mm) (Freudenberg Medical Europe GmbH) was manufactured from medical grade silicone elastomer (MED-4050 by NuSil Technology) for the body. Considering that the tube slightly expands when filled with the silicone elastomer, an overall diameter of approximately 0.95 mm was expected.

The main technological challenges for the implant electrodes are charge-injection capacity, reliability and biocompatibility. Pt-Ir electrodes are widely used because they combine the good charge-injection capacity of iridium with the durability and ease of manufacturing of platinum [38]. The helically coiled wire electrode configuration provides stretchability, a larger surface area to decrease the current density without increasing the overall size and offers a surface that can promote good anchorage to reduce implant dislocation without causing tissue damage [38–40].

Each electrode and the interconnections to the feedthroughs were manufactured as a single coil to avoid additional connection points. This was possible by manufacturing coils with variable pitch and diameter. These coils define the two electrodes located at opposite ends of the implant body, and the two interconnections (0.6 mm diameter, 10.25 mm length) to the feedthrough pins of the capsule. The length of the interconnection coil was set to achieve the established separation between the electrodes (**Figure 6**(e)). The interconnection coils were electrically connected to the feedthrough pins using laser welding to avoid the combination of different materials that might produce corrosion [36]. The reduced dimensions of all the parts and components of the implantable microstimulator before the final implementation can be observed in **Figure 7**. Using a silicone swelling fluid (Opteon™ SF79 by The Chemours Company FC, LLC), the structure was then introduced in the custom-made silicone tube, leaving only the electrodes exposed. A piece of silicone tube was inserted to support the electrodes and as an entry point to fill the tube with a medical grade silicone elastomer (MED-6015 by NuSil Technology), securing the whole assembly after curing (**Figure 6**(f)).

**Figure 7.**
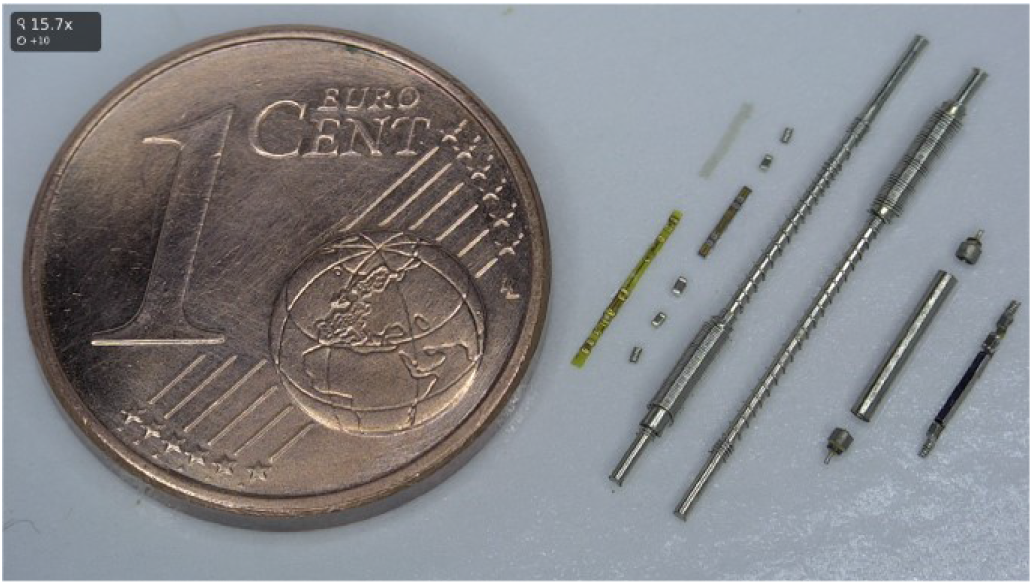
Parts and components of the electronic assembly, electrodes, and hermetic capsule before implementation.

### 2.2 In vivo study

#### 2.2.1 System modeling

As previously demonstrated in [30,41–43], volume conduction can be electrically modeled using a two-port impedance network that characterizes coupling between the external system and the implanted device. Here, such modeling approach was tried for further confirmation of its validity. In particular, it was assayed its use for modeling the behavior of the ASIC in the animal assays carried out in this study. For that, the two-port impedance model was combined with the SPICE model of the ASIC.

Here, as in [30], the electrode impedances were neglected considering they are low compared to tissue impedances. Tissue impedances were approximated as resistances, which were calculated by imposing voltages and currents across the external and the implant electrodes in simulations. See supplementary material for details on the two-port network calculations.

The three impedances of two-port impedance model [42] were obtained for a geometry (**Figure 8**) modelling the small animal model assayed in this study. These parameters were obtained by performing a finite element method (FEM) simulation in COMSOL Multiphysics® 5.3 using the “Electric Currents” physics interface in the AC/DC module. The geometrical model approximated the anatomy of the rabbit hind limb (**Figure 8**) obtained from measurements on the x-ray images taken during the animal procedures (including bone, muscle and skin). The geometries of the epidermal electrodes and the implant were added to the model according to the materials and setup used in the *in vivo* animal study described below. The passive dielectric properties used for the simulation were established for a frequency of 3 MHz (Table I of supplementary material), which is the frequency used in the animal study. The FEM simulation software automatically generated a mesh of 106933 tetrahedral elements.

**Figure 8.**
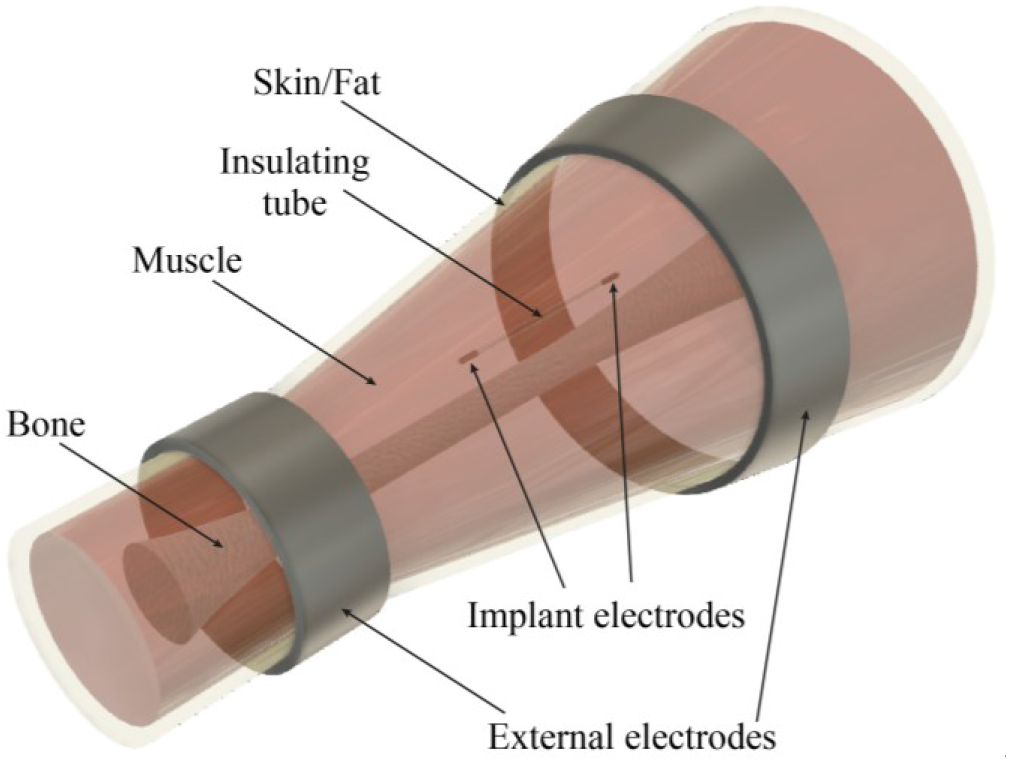
Geometry used to create a two-port impedance network for modelling volume conduction coupling between the external system and the implant electronics in the animal model assayed in this study. (See text).

#### 2.2.2 External generator

The architecture of the external generator used to perform the experiments described here is shown in Figure 9. This system allows to address multiple microstimulators and to define the stimulation parameters (current amplitude, pulse repetition frequency and pulse duration).

**Figure 9.**
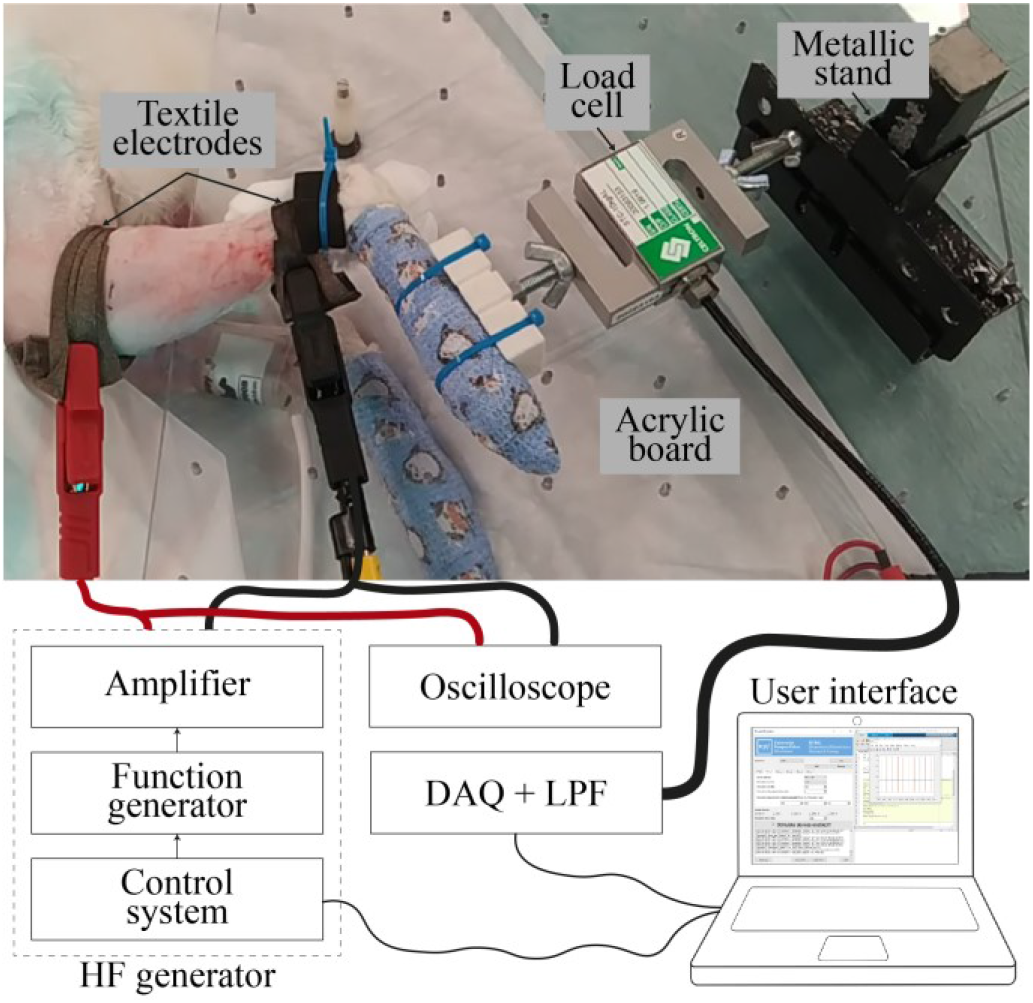
External generator and force measurement setup for *in vivo* experimentation. The ankle of the animal was fixed to an acrylic board and the foot was fixed to a load cell to measure dorsiflexion and plantarflexion isometric contraction forces. The textile electrodes were strapped around the limb and were connected to the external generator.

The system consists of a computer, a digital controller, a function generator and an amplifier. A Python-based user interface running in the computer sets the parameters of the digital controller via serial port. The controller is custom-made and FPGA-based. It generates a modulating signal for the function generator. The function generator (AFG 3022B by Tektronix Inc.) uses this signal to modulate a 3 MHz sinusoidal voltage carrier using amplitude-shift keying (ASK) modulation (OOK: On–off keying). The modulated waveform is then amplified using a custom-made high voltage amplifier [30] and delivered across the tissue through two external electrodes. For the in vivo demonstration, the external electrodes consisted of 1.5 cm wide bands made from silver-based stretchable conductive fabric (Shieldex® Technik-tex P130 + B by Statex Produktions und Vertriebs GmbH) strapped around the target limb of the animal.

#### 2.2.3 Force measurements

The force measurement setup is also shown in **Figure 9**. Isometric plantarflexion and dorsiflexion forces evoked during stimulation were measured using a S-type load cell (Celtron STC-5kgAL by Vishay Precision Group, Inc.). A custom-made acrylic board was used for securing with bolts a metallic stand to which the load cell was also secured with bolts. With the animal lying sideways, the limb to be tested was placed on top of the acrylic board, flexing the ankle joint at approximately 90° to optimize muscle length and response to electrical stimulation. The limb was fastened at the ankle using cable ties and an atraumatic pad. The foot was protected by applying self-adherent bandaging tape and cable ties were used to fasten the foot to a plastic plate attached to the load cell. The force signals from the load cell were acquired using a custom-made signal conditioning system composed of a precision instrumentation amplifier with a voltage gain of 100 V/V (LT1101 by Analog Devices, Inc.), a RC low-pass filter (LPF) (cut-off frequency: 2 kHz), and an isolation amplifier with a fixed gain of 8.2 V/V (ACPL-790B by Broadcom, Corp.), followed by a data acquisition (DAQ) board (NI-USB6216 by National Instruments Corp.) configured at a sampling rate of 25 kHz. These signals were digitally filtered using a third-order Butterworth LPF (cut-off frequency: 10 Hz) in MATLAB® (by The MathWorks, Inc.).

#### 2.2.4 Electrical measurements

Electrical measurements of voltages and currents applied through the external electrodes in the *in vivo* setup were performed and recorded using an active differential probe (TA043 by Pico Technology Ltd.), a current probe (TCP2020A by Tektronix, Inc.) and a battery-powered oscilloscope (TPS2014 by Tektronix Inc.).

To explore the electrical behavior of the implant, a bipolar probe and a test circuit were also used for measurement purposes. The probe consisted in a 1.17 mm diameter coaxial cable (Filotex® ET087059 by Nexans S.A.) whose core and shield conductor were used as electrodes (**Figure 10**). The insulation and shield of the cable were removed from the tip of the cable, leaving 3 mm of the core wire exposed, and a 3 mm section of the outer insulation was removed at 3 cm from the tip, leaving the shield exposed. Then, a 1.3 mm-diameter stainless-steel ring was placed in contact with the shield conductor. By doing so, the bipolar probe electrodes are similar to those of the implantable stimulators.

**Figure 10.**
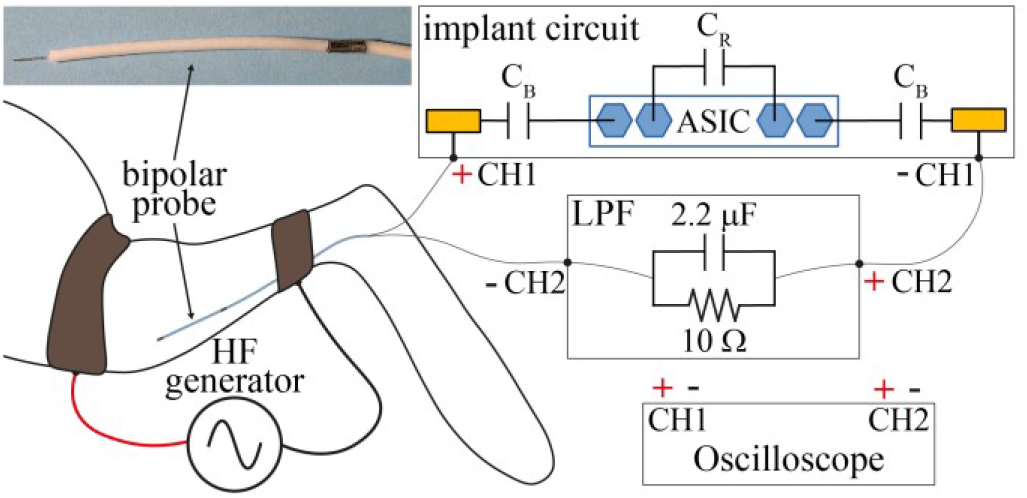
Electrical measurements setup for the *in vivo* experimentation. Before the implantation of the microstimulator, a bipolar probe with electrodes similar to those of the implant was implanted in the same location and connected in series to an external implant circuit and a low-pass filter (LPF) to measure and record the voltage across the implant electrodes and the low frequency current flowing through the device.

The test circuit consisted in a non-encapsulated implant circuit of the microstimulators demonstrated here. Also, a firs-order LPF with a cut-off frequency of 7.2 kHz (10 Ω resistor and 2.2 μF capacitor connected in parallel) was used to obtain the LF current. The setup is shown in **Figure 10**. The test circuit was connected in series to the probe and to the LPF for measuring and recording the voltage difference across the implant electrodes and the LF current flowing through the device using the oscilloscope.

#### 2.2.5 Animal handling

The *in vivo* animal study was approved by the local Ethical Committee for Animal Research of the Barcelona Biomedical Research Park (CEEA-PRBB), and by the Catalan Government (application number: BPM-18-0028AE, project number: 10109). Two New Zealand White male rabbits, weighting approximately 6 kg, were employed for acute implantation of the microstimulators and stimulation assays procedures.

For sedation and initial anesthesia, dexmedetomidine (50 μg/kg), butorphanol (0.4 mg/kg) and ketamine (10 mg/kg) were intramuscularly administered prior to the preparation of each animal. In each case, both hind limbs of the rabbit were shaved, from the head of the femur to the mid tarsus and a depilatory cream (Veet sensitive skin by Reckitt Benckiser Group plc) was applied for 1.5 minutes and thoroughly rinsed for complete hair removal in the specified areas. Finally, the rabbit was transferred to an anesthetic circuit using endotracheal intubation and anesthesia was maintained with isoflurane (2-3 %). During the entire procedure, a maintenance fluid with 5 % glucose-saline (8 mL/h) was administered intravenously, and a heating pad was employed. The animals were constantly monitored via capnography, pulse oximetry and esophageal temperature measurement. At the end of the session with each rabbit, the animal was euthanized by intravenous injection of sodium pentobarbital (10 mL).

#### 2.2.6 Implantation procedure

Two muscles in the rabbit hind limb were selected as target muscles and identified by palpation: tibialis anterior (TA), and gastrocnemius lateralis (GL). The procedure followed for the implantation of the microstimulators is depicted in **Figure 11**. A 14 G (2.1 mm outer diameter, 1.529 mm inner diameter, 83 mm length) catheter (BD Angiocath 382268 by Becton, Dickinson and Company) was introduced longitudinally in the target muscle (**Figure 11**(a)). Then, the tip of the catheter needle was used as an exploration electrode to verify that the location was suitable for stimulation: a commercial current-controlled electrical stimulator (Kegel8®) was used to deliver symmetric biphasic pulses of 2 mA, 200 μs duration, at 100 Hz, across the catheter needle and an Ag/AgCl gel electrode (Red Dot™ 2239 by 3M) placed on the thigh of the animal as return electrode. The catheter tip was repositioned if required to ensure that the induced contraction was sufficiently strong for observation. Once the location of the catheter tip was deemed satisfactory, the catheter needle was removed (**Figure 11**(b)) and the implantable microstimulator was introduced into the catheter tube (**Figure 11**(c)), ensuring electrode A was facing the tip, and it was pushed towards the distal end at the motor point using a custom-made pusher (**Figure 11**(d)). At this point, the catheter tube was carefully retracted while holding the pusher in place to release the implantable device within the muscle (**Figure 11**(e)). Finally, both the pusher and the catheter were completely removed (**Figure 11**(f)).

**Figure 11.**
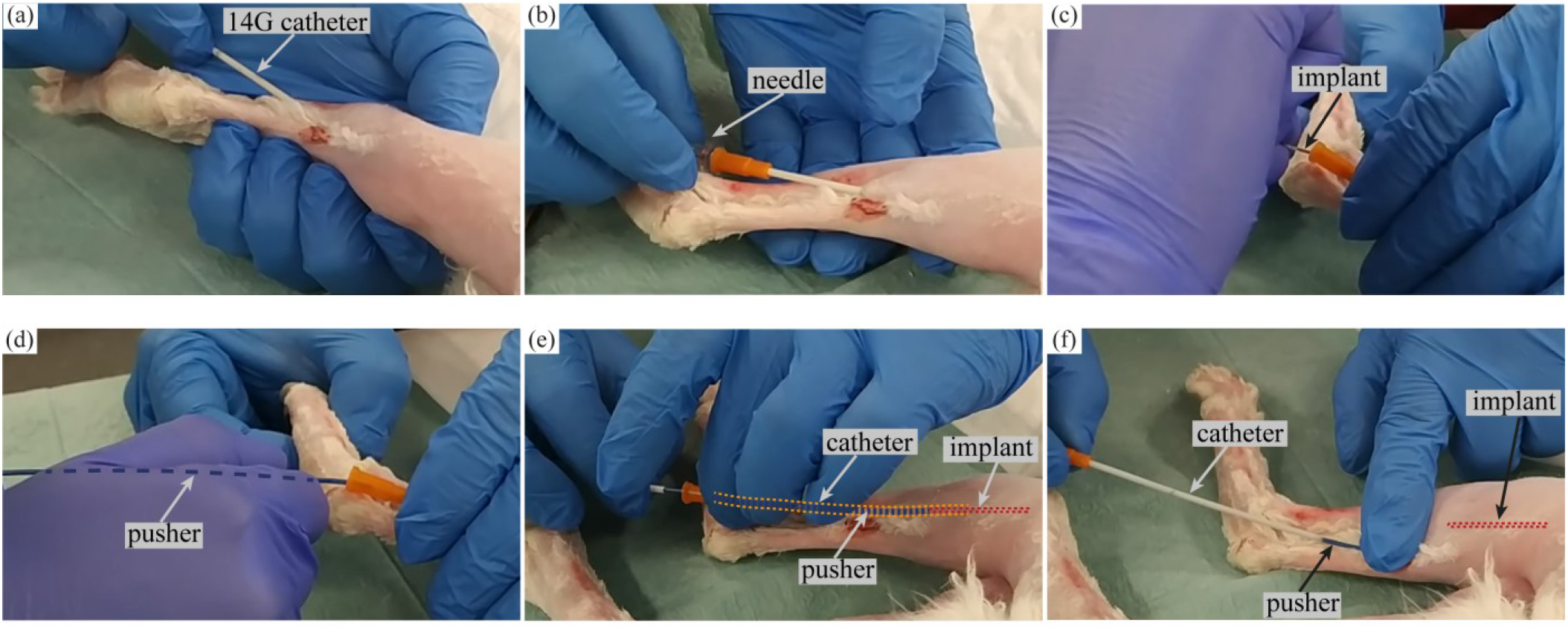
Implantation procedure. (a) A 14 G catheter was introduced in the target muscle. (b) The catheter needle was removed from the outer tube. (c) The implant was inserted into the catheter tube. (d) A pusher was used to push the implant towards the end of the catheter tube, leaving the stimulation electrode of the implant (electrode A) at the motor point. (e) Holding the pusher in place, the catheter tube was retracted to release the implant within the muscle. (f) The pusher and the catheter tube are fully removed.

On a few occasions, before introducing the implant through the catheter, the previously described bipolar probe was introduced through the catheter and was externally connected to the test circuit to perform the indicated electrical measurements (**Figure 10**). For that, the catheter had to be retracted 3 cm for exposing the proximal electrode of the bipolar probe. After concluding the measurements, the catheter was pushed back 3 cm, the bipolar probe was removed, and the implant was introduced and implanted as described in the above paragraph.

#### 2.2.7 Stimulation assays

A pilot session with the first rabbit was performed to assay the implantation procedure and to fine-tune the external generator and measurements setup. In the following session with the second rabbit, a series of stimulation assays were performed to carry out electrical and force measurements. The stimulation sequences consisted of several HF bursts addressed at the same implanted microstimulator and were delivered at a constant repetition frequency (*F*). Single-device stimulation sequences had a burst duration *B*) of 300 ms, and multiple-device stimulation assays had a duration of 400 ms.

## 3. Results

### 3.1 ASIC in vitro assays

The ASIC design submitted to the foundry contained seven ASICs to be sub-diced: 6 final form factor circuits (with different hard-wired addresses) and an additional wider prototype that included extra bonding pads for characterization and checking crucial nodes (analog and digital). In the final form factor circuits (i.e., those of the implants), the only available bonding pads are the two connections to the dc-blocking capacitors (C_B_) and the external reservoir capacitor (C_R_) connections (**Figure 12**). The device was first thoroughly characterized electrically (results not reported here) before performing *in vitro* and *in vivo* tests.

**Figure 12.**
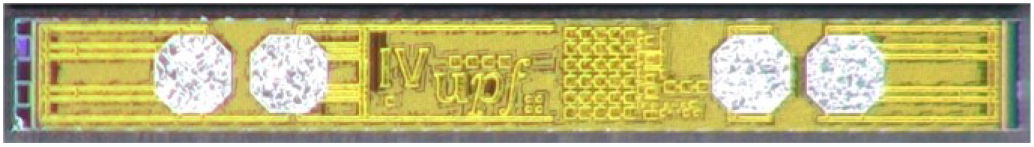
Picture of the ASIC, after sub-dicing, with the four bonding pads for external connections. Its final dimensions are approximately: 300 μm x 2.5 mm x 204 μm

After electrical characterization, the ASICs were tested in an *in vitro* test setup as depicted in **Figure 13**. A saline agar cylinder (14 cm long, 7 cm diameter) with a conductivity of about 0.57 S/m was used to model muscle tissue. Two wire electrodes with an area roughly equivalent to that of the implant electrodes (0.25 mm diameter single wire conductor, exposed 6 mm; bent 180° in the middle) were introduced in the saline agar cylinder and connected as depicted in **Figure** 13. A HF signal with a voltage amplitude of 20 V was applied through two aluminum band electrodes (width = 10 mm, separation distance = 8 cm).

**Figure 13.**
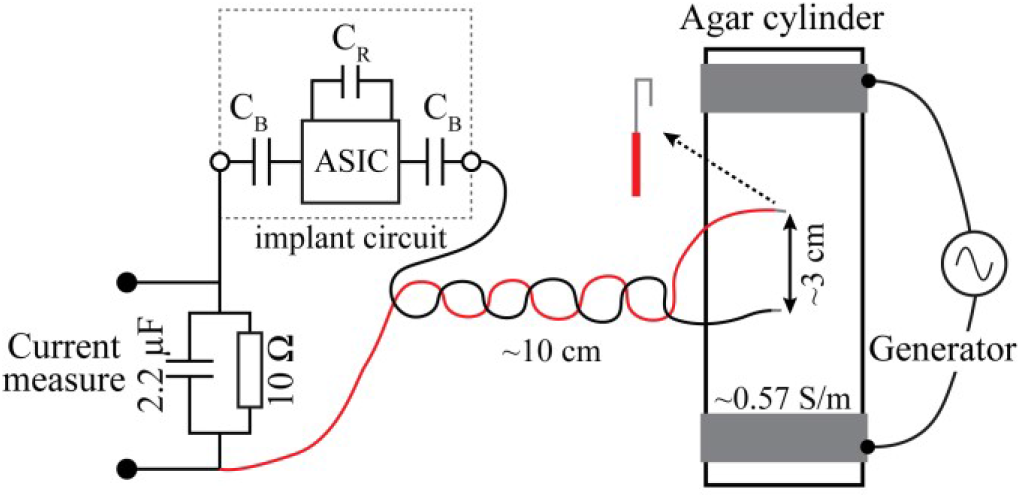
Test setup used for *in vitro* measurements.

The ASICs were correctly addressed to selectively trigger flow of rectified current and the current amplitude could be selected (**Figure 14)**.

**Figure 14.**
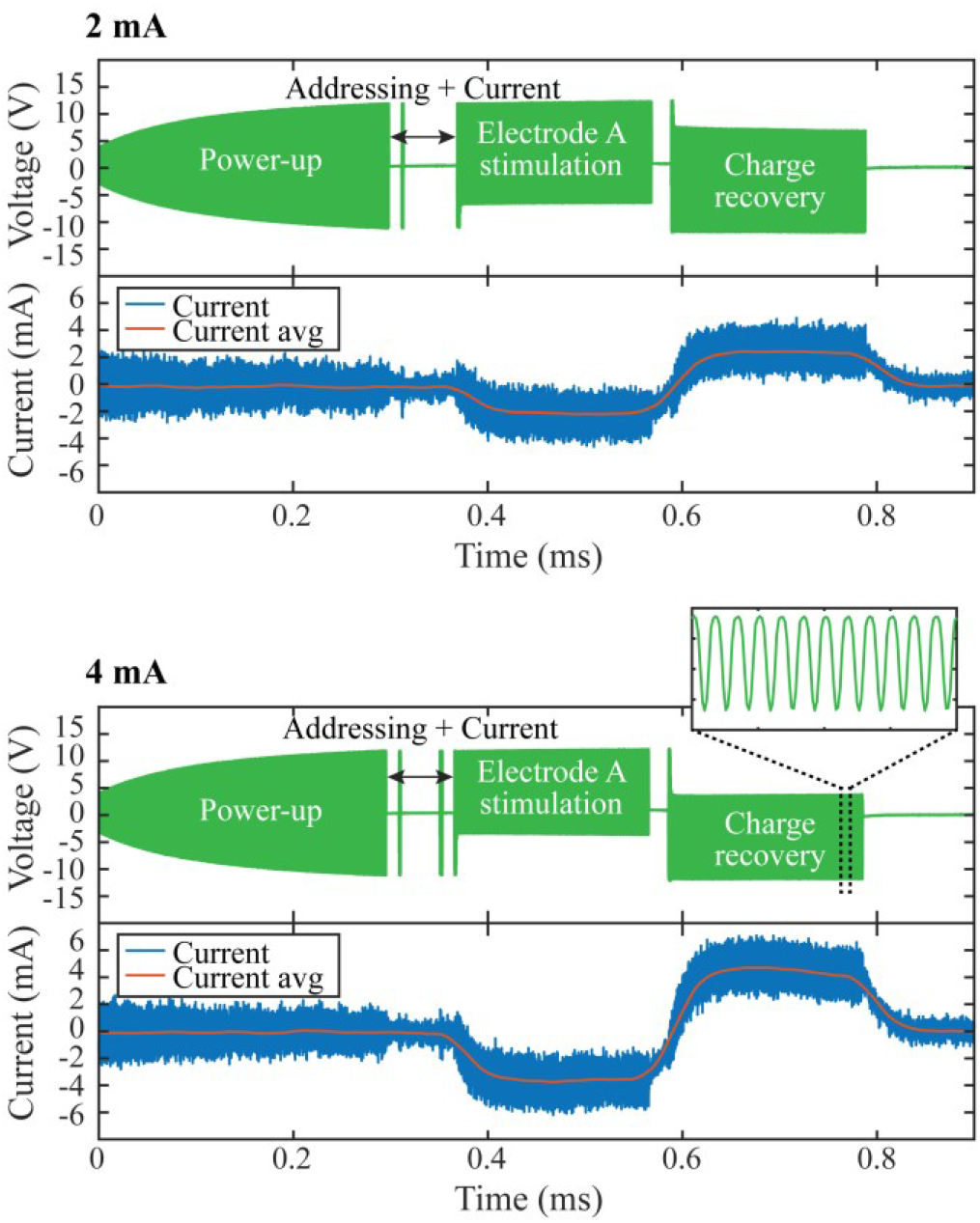
Voltage across (green) and low frequency current through (blue) the implant circuit during two *in vitro* assays. The red trace is a smoothed version (rolling average of 1250 samples, sampling frequency = 31.25 MHz) of the low frequency current. The same implant circuit was used in both assays (address = 4).

### 3.2 In vivo study

The first implantation procedure was performed into the TA muscle in the hind limb of the rabbit.

Before implanting the microstimulator, the above referred bipolar probe was deployed through the catheter and was externally connected to the test circuit to perform electrical measurements (**Figure 10**). By delivering through the textile electrodes (minimum distance between electrodes = 6 cm) a HF burst signal (*f* = 3 MHz, *V_ext_* = 42 V, *F* = 100 Hz, *B* = 300 ms, power-up = 300 μs, stimulation = 200 μs, gap = 20 μs, *I_implant_* = 2 mA), stimulation was achieved only when the address corresponding to the test circuit (address 6) was configured in the addressing stage of the signal. The measured current amplitude of the delivered HF signal was 177 mA. A voltage amplitude of 4.76 V was measured across the probe electrodes. **Figure 15** shows the recorded LF current. The recorded signal indicates that the implant was able to apply biphasic LF currents with the configured parameters.

**Figure 15.**
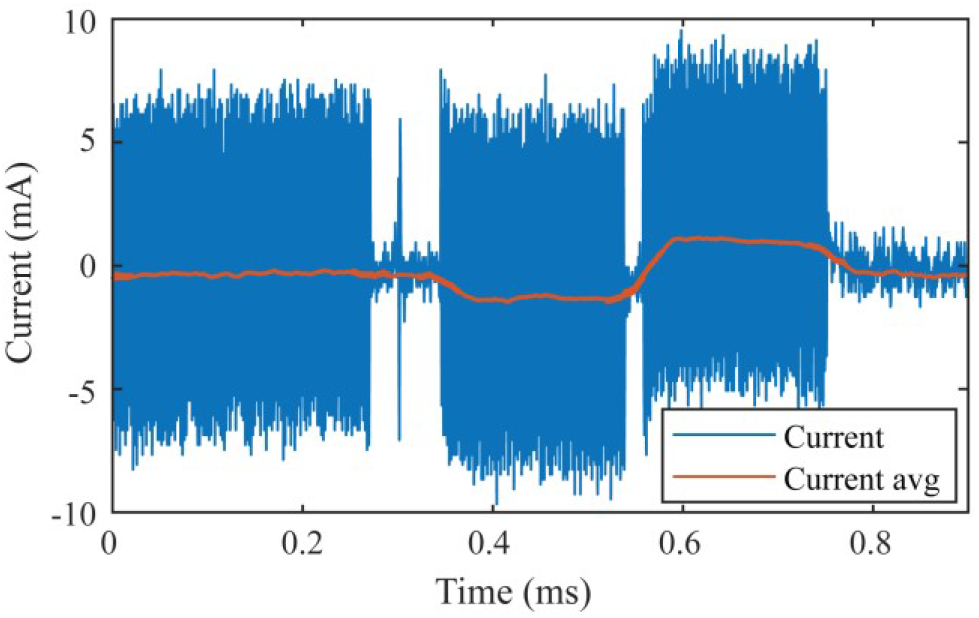
Measured low frequency current through the bipolar probe connected to the implant test circuit during the first *in vivo* assay. The red trace is a smoothed version (rolling average of 100 samples, sampling frequency = 2.5 MHz) of the low frequency current.

As indicated in section 2.2.1, the two-port network parameters obtained for this animal model were included in a SPICE simulation of the ASIC. Setting the amplitude voltage across the external electrodes (*V_ext_* = 42 V) in the simulation, a voltage amplitude of 4.91 V was obtained across the implant electrodes. This result reinforces the validity of the modeling approach.

After the implantation of the first device (address 1) in the same location in the TA muscle (**Figure 16**(a)), stimulation assays were performed. The first reported stimulation sequence was performed by delivering across the external electrodes a HF signal (*f* = 3 MHz, *V_ext_* = 41 V, *B* = 300 ms, power-up = 300 μs, stimulation = 200 μs, gap = 20 μs, *I_implant_* = 2 mA) starting with a repetition frequency (*F*) of 20 Hz and increasing it to 50, 100 and 200 Hz. The measured current amplitude of the HF signal was 200 mA. **Figure 16**(b) shows that the force magnitude exerted by the foot attached to the load cell increases with the frequency of the bursts between 20 and 100 Hz, confirming that force modulation was possible by varying the repetition frequency of the stimulation signal, although, as expected, minor differences were observed between 100 and 200 Hz. A video of this stimulation sequence is available as supplementary material.

**Figure 16.**
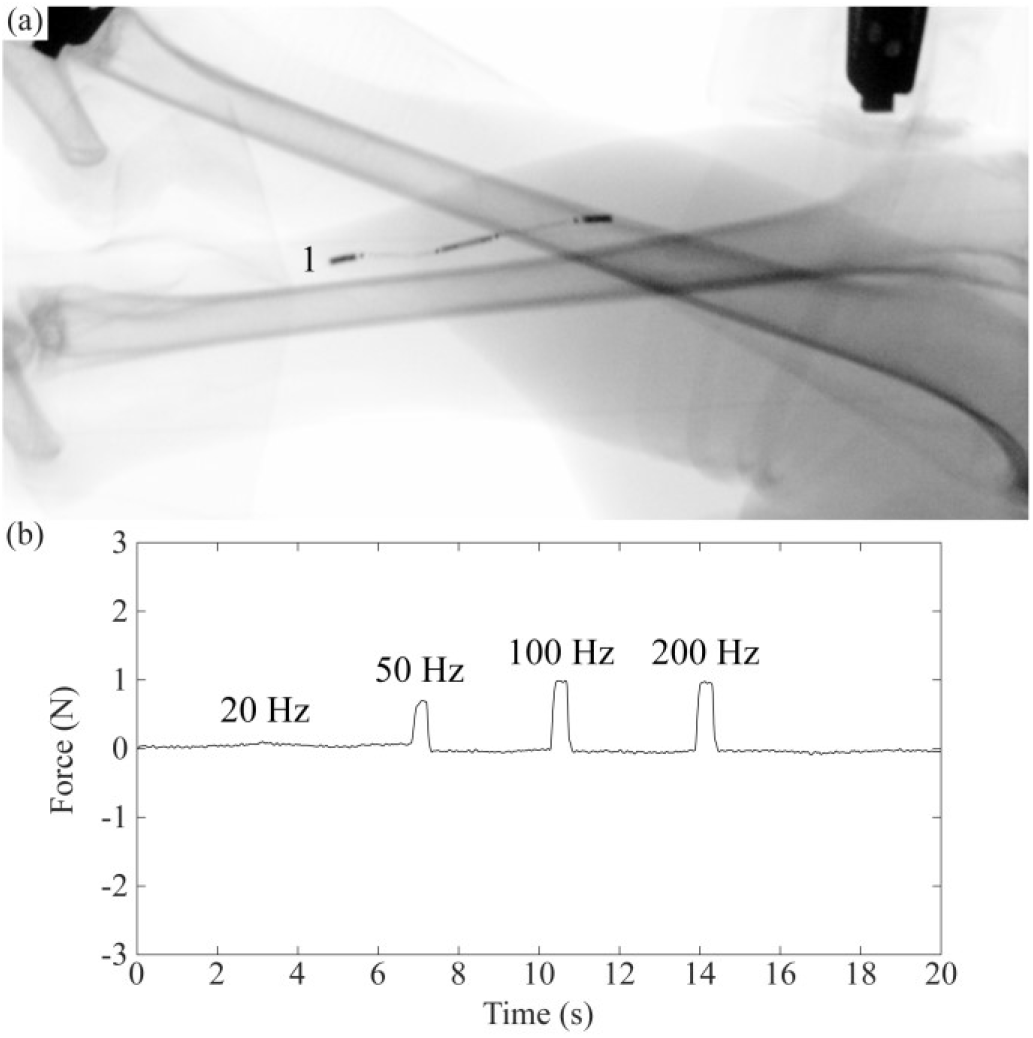
Implantations and isometric force measurements. (a) x-ray image of first implant deployed into the tibialis anterior (TA) muscle. (b) Force recordings. The repetition frequency (*F*) of the HF bursts (*f* = 3 MHz, *V_ext_* = 41 V, *B* = 300 ms, power-up = 300 μs, stimulation = 200 μs, gap = 20 μs, *I_implant_* = 2 mA) delivered across the external electrodes was increased from 20 Hz to 200 Hz to modulate the force magnitude.

A second device (address 2) was implanted also in the TA muscle and a third device (address 3) was implanted in the GL muscle **Figure 17**(a). After each implant was deployed, further stimulation assays were performed. In the second reported stimulation sequence, these three devices were powered and individually addressed delivering HF bursts (*f* = 3 MHz, *V_ext_* = 49 V, *F* = 100 Hz, *B* = 400 ms, power-up = 300 μs, stimulation = 200 μs, gap = 20 μs, *I_implant_* = 2 mA) through the external system. In this sequence, the measured current amplitude of the delivered HF signal was 384 mA. A video of this stimulation sequence is available as supplementary material.

**Figure 17.**
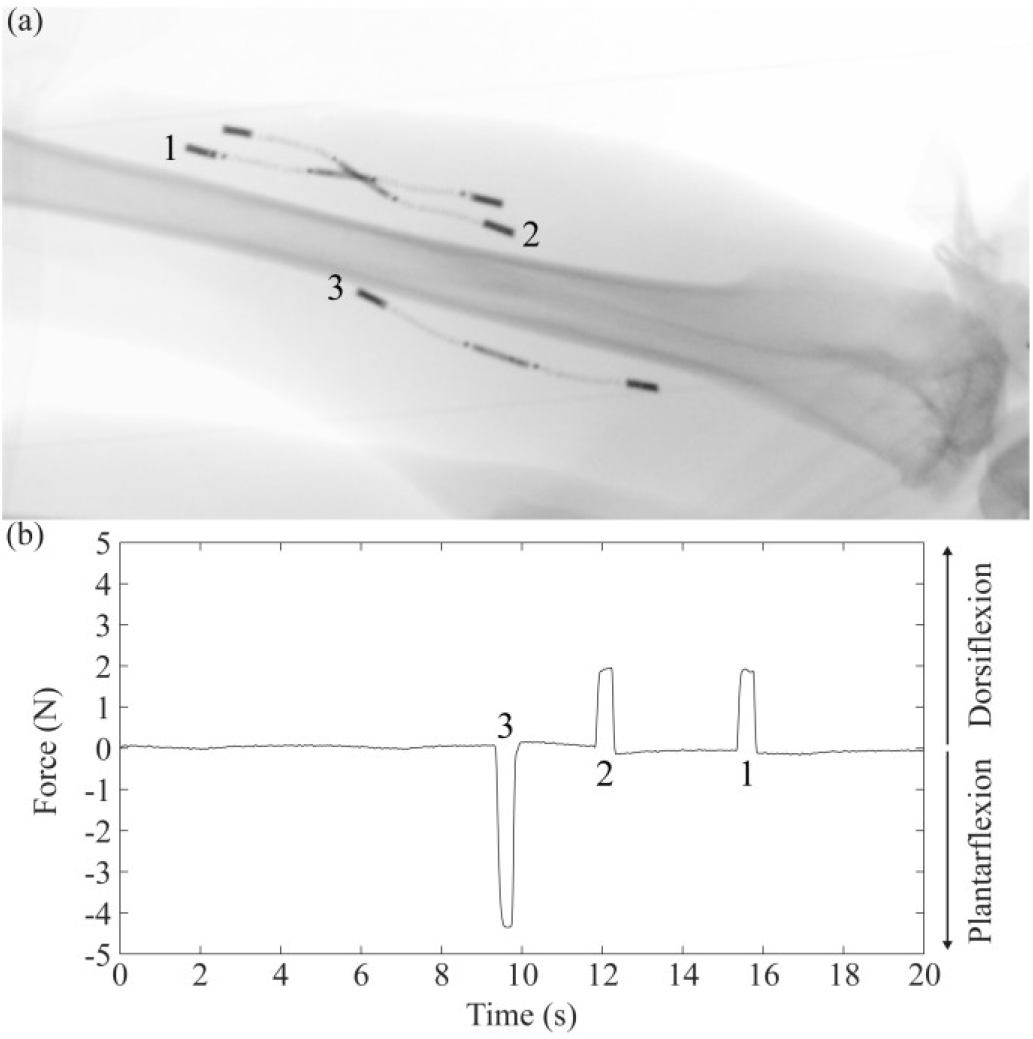
Implantation and isometric force measurements. (a) x-ray image of three implants deployed by injection, 1 and 2 into the tibialis anterior (TA) and 3 into the gastrocnemius lateralis (GL) muscles. (b) Force recordings. The implants were independently addressed to induce plantarflexion or dorsiflexion movements by delivering HF bursts (*f* = 3 MHz, *V_ext_* = 49 V, *F* = 100 Hz, *B* = 400 ms, power-up = 300 μs, stimulation = 200 μs, gap = 20 μs, *l_implant_* = 2 mA) across the external electrodes.

## 4. Discussion

The injectable microstimulators *in vivo* demonstrated here have been built using fabrication techniques and materials well established for chronic electronic implants. This ensures that major regulatory roadblocks will be avoided, thus paving the way to the clinical development of neuroprostheses formed by these devices. This contrasts with other developments targeting implant miniaturization that do not address issues such as the proven long-term biocompatibility of the materials or the regulatory acceptance of the electronic packaging methods [44].

Because of their thinness, flexibility, and the minimal invasiveness of their implantation procedure, we anticipate it will be possible to deploy these implants in large quantities, hence allowing the development of advanced motor neuroprostheses formed by dense networks of such wireless devices. For instance, we envision their use to perform multi-site interleaved intramuscular stimulation to reduce muscle fatigue and smooth forces for movement restoration [45].

A key innovation of the present study has been the development of an integrated circuit that acts as a “smart rectifier”. To the best of our knowledge, this is the first addressable integrated circuit capable of performing stimulation based on rectification of HF currents delivered by volume conduction. The integrated circuit is not only addressable but it also allows to: 1) define the sense of the rectified current (and hence the sense of the stimulation current), 2) limit the magnitude of the rectified current regardless of the local magnitude of the HF electric field, and 3) select the limit for the magnitude of the rectified current (only two limits were demonstrated here but it is obvious that the circuit can be upgraded to allow many more limits and, in fact, we are already working on such upgrade). As indicated above, further details of the design of the integrated circuit and its characterization, as well as a discussion of its major challenges will be presented in a subsequent publication. Nevertheless, here there are other innovations and aspects worth discussing in some detail:

### 4.1 ASIC bonding innovation

The combined use of conducting polymers and compression force applied by thermally shrinkable tubing for creating permanent electrical connections to integrated circuit bond pads is, to the best of our knowledge, unprecedented. The anisotropic conducting polymer used for the electronic assembly of the microstimulators is intended for integrated circuits and subassemblies testing applications in which several hundred thousand contact cycles are expected. Loosing or removal of the surface conducting particles is the main failure mechanism. Given that this mechanism is not present in this application, high durability is anticipated. The pads surface area should contact a sufficient number of the conducting columns in the polymer to avoid a considerable series resistance [46]. The version of the material used is designed for PCB pads of at least 0.06 mm in diameter and a combined pad height of 0.018 mm, which is suitable for the pad dimensions of the PCB included as a substrate and for the ASIC pads (**Figure 6**(c)).

### 4.2 Hermetic capsule

The use of the thin titanium capsule sealed by laser welding, with feedthroughs manufactured with brazing technology for insulated electrical connections, provides hermeticity and, assumably, improves biocompatibility and mechanical stability with respect to previous prototypes [36,37,47]. A noteworthy feature of the cylindrical capsule design is that it offers flexibility to easily adapt its length for different requirements of the ASIC and other electronic components.

Although titanium encapsulation is known to provide superior hermeticity [36] and the feedthroughs have been manufactured to be hermetic, in truth, the hermeticity of the capsule has not been demonstrated here. Standardized methods for hermeticity characterization (EN ISO 20485:2018: Tracer gas method [48] and MIL-STD-883-1, Method 1014.17: Seal [49]) basically consist in pumping in and sensing the amount of a tracer gas leaking through the materials or sealings. There are limitations for using these gas leakage methods to test encapsulations and components with cavities of volumes below 10^-3^ cm^3^, considering that the resolution and detection mechanisms available are incompatible with such small volumes. The cavity of the hermetic capsule presented here has a volume of approximately 7 × 10^-4^ cm^3^, which is below the restrictions of the standard methods. This suggests that test methods have to be refined and improved for the adequate hermeticity characterization of the devices described here [50,51].

### 4.3 Implant body thinness

The thin threadlike flexible body of the implantable microstimulators, with a maximum diameter of 0.97 mm, allows to perform minimally invasive implantation procedures. This has been demonstrated in the *in vivo* animal study where the implantation was performed by injection using a 14 G catheter. The minimum inner diameter of this catheter is 1.529 mm, which is considerably wide for the implant dimensions. The following gauge commercially available for the catheter used is 16 G, with a minimum inner diameter of 1.125 mm. The implantation procedure was attempted with this catheter, however, the rubber-like surface of the silicone body difficulted the retraction of the catheter tube. A custom-made catheter with a fitted diameter and a coating adequate for the body surface of the device would further improve the implantation procedure.

### 4.4 Electrical safety

Compliance with safety standards for human exposure to electromagnetic fields defined by IEEE [52] and the International Commission on Non-Ionizing Radiation Protection (ICNIRP) [53] ensures innocuousness and thus facilitates approval by medical devices regulatory bodies. Based on both standards, the limits to avoid the risk of thermal damage due to Joule heating are established using the Specific Absorption Rate (SAR). The SAR indicates the heat dissipated per unit of tissue mass and is calculated averaging over 10 g of mass and 6 minutes. Previous studies have demonstrated, numerically [54] and in humans [30], that it is possible to transfer powers in the order of milliwatts to implanted devices via volume conduction of HF current bursts that comply with these safety standards.

In the present study, as an addendum, it was verified that the developed microstimulators can operate with SAR levels compliant with the standards. For that, the minimum amplitude of the HF current required for implant operation was measured in an *in vitro* setup similar to that described for the ASIC *in vitro* tests. Stimulation currents were produced when a current amplitude of 0.68 A (*I_ext_*) was flowing through the cylindrical saline agar phantom (NaCl 0.3%; conductivity, *σ*, = 0.52 S/m) with a diameter of 5 cm. Thus, the current density around the implant location can be approximated as

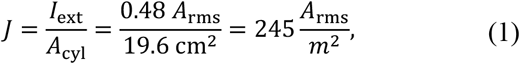

where *A_cyl_* is the area of the cylinder.

Applying the Ohm’s law,

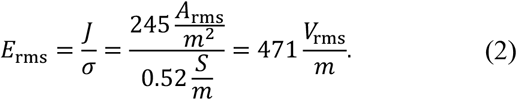

The delivered HF signal (power-up = 300 μs, stimulation = 200 μs, gap = 20 μs) had a burst duration *B* = 700 μs and a repetition frequency F = 100 Hz. Therefore, its duty cycle was

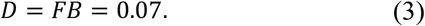

Consequently, the average electric field was

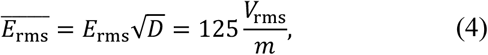

and the SAR was

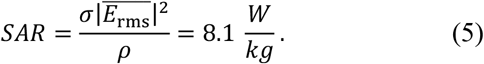

This SAR, which was calculated for an implausible extreme scenario (continuous neuromuscular stimulation at 100 Hz), is below the limit of 10 W/kg dictated by the standards (20 W/kg for limbs). This demonstrates that the microstimulators presented here can be operated by delivering HF currents that comply with the SAR limits. Nevertheless, it is worth noting that, if needed, substantial improvement in this regard could be easily obtained by elongating the implants [24].

## 5. Conclusion

Threadlike microelectronic stimulators, capable of picking up HF currents delivered by volume conduction and converting them into LF currents that can electrically stimulate excitable tissues, have been developed and demonstrated. These devices have been built using fabrication techniques and materials well established for chronic electronic implants. Their submillimetric dimensions and flexibility make them suitable for minimally invasive implantation by injection. These devices are addressable, allowing to establish a network of microstimulators that can be controlled independently. Their operation was demonstrated by intramuscularly implanting a series of these devices in the hind limb of anesthetized rabbits, using a 14 G catheter, and inducing controlled and independent muscle contractions. We believe this study paves the way to the clinical development of a next generation of motor neuroprostheses formed by dense networks of such wireless devices.

## Acknowledgments

The authors would like to express their gratitude to Nerea Álvarez de Eulate Llano and Jaume Millán i Ichon (technical support staff at UPF) for their assistance in the manufacturing of the implants, and to Jordi Grífols and all the team at the Centre de Medicina Comparatives i Bioimatge de Catalunya (CMCiB) for their work regarding all animal procedures.

