## Supplementary information for "Wireless networks of injectable microelectronic stimulators based on rectification of volume conducted high frequency currents"

**Supplementary material**

### Methods:

#### System modeling

As previously demonstrated in [1]–[4], volume conduction can be modeled using a two-port impedance network that characterizes coupling between the HF generator and the implantable device. This network models the impedance of skin and implant electrodes, and the tissue impedances coupled between them (**Figure 1**). Voltages and currents at this network can be expressed as

$$\begin{bmatrix} v_1 \\ v_2 \end{bmatrix} = \begin{bmatrix} Z_{11} & Z_{12} \\ Z_{12} & Z_{22} \end{bmatrix} \begin{bmatrix} i_1 \\ i_2 \end{bmatrix}, \qquad (1)$$

where $v_1$ is the voltage across the epidermal electrodes and $v_2$ is the voltage across the implant electrodes, $i_1$ is the current through the epidermal electrodes, $i_2$ is the current through the implant and $z_{xy}$ are the impedance parameters [2].

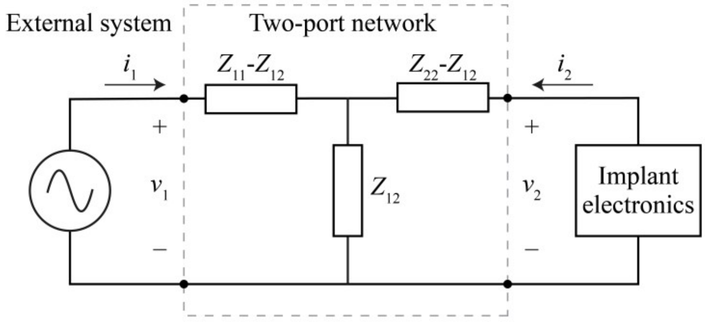


**Figure 1**. Two-port network modelling the coupling between the external system and the implantable device.

Considering that electrode impedances are low compared to the remaining system impedances and that tissue impedances can be approximated as resistances, the impedances in (1) can be substituted by resistances, resulting in

$$\begin{bmatrix} V_1 \\ V_2 \end{bmatrix} = \begin{bmatrix} R_{11} & R_{12} \\ R_{12} & R_{22} \end{bmatrix} \begin{bmatrix} I_1 \\ I_2 \end{bmatrix} \qquad (2)$$

By definition

$$R_{11} \stackrel{\text{def}}{=} \left.\frac{V_1}{I_1}\right|_{I_2=0}, R_{22} \stackrel{\text{def}}{=} \left.\frac{V_2}{I_2}\right|_{I_1=0}, R_{12} \stackrel{\text{def}}{=} \left.\frac{V_1}{I_2}\right|_{I_1=0} \qquad (3)$$

and, by imposing these conditions, $R_{11}$, $R_{22}$, and $R_{12}$ can be obtained using numerical modelling. Performing a FEM simulation in COMSOL Multiphysics® 5.3 of a model based on an approximation of the anatomy of the rabbit hind limb using electrical parameters for a frequency of 3 MHz (Table I), $R_{11}$and $R_{12}$ were calculated by applying a known electric current through the epidermal electrodes and measuring the voltage across the epidermal electrodes and the implant electrodes. In a similar manner, $R_{22}$ and $R_{21}$ were calculated by applying the current through the implant electrodes. The resulting values of these parameters are: $R_{11} = 154.2$, $R_{22} = 331.85$, and $R_{12} = 35.21$.

TABLE I
DIELECTRIC PROPERTIES OF MATERIALS AT 3 MHZ

| Material | Conductivity σ (S/m) | Relative permittivity |
|---|---|---|
| Bone (cortical) [5] | 0.032 | 83 |
| Muscle [5] | 0.57 | 5220 |
| Skin/fat [5] | 0.052 | 303 |
| Implant electrodes [6] | $4\times10^{6}$ | 1.6 |
| Insulating tube [7][8] | $1\times10^{-12}$ | 4.3 |
| External electrodes [9] | $4.4\times10^{7}$ | 1.6 |
